# Laminar Segregation of Sensory Coding and Behavioral Readout in Macaque V4

**DOI:** 10.1101/399626

**Authors:** Warren W. Pettine, Nicholas A. Steinmetz, Tirin Moore

## Abstract

Neurons in sensory areas of the neocortex are known to represent information both about sensory stimuli and behavioral state, but how these two disparate signals are integrated across cortical layers is poorly understood. To study this issue, we measured the coding of visual stimulus orientation and of behavioral state by neurons within superficial and deep layers of area V4 in monkeys while they covertly attended or prepared eye movements to visual stimuli. We show that single neurons and neuronal populations in superficial layers convey more information about the orientation of visual stimuli, whereas single neurons and neuronal populations in deep layers convey greater information about the behavioral relevance of those stimuli. In particular, deep layer neurons encode greater information about the direction of prepared eye movements. These results reveal a division of labor between laminae in the coding of visual input and visually guided behavior.

## Introduction

Visual area V4 comprises an intermediate processing stage in the primate visual hierarchy^1,2^. V4 neurons exhibit selectivity to color^3,4^, orientation^5,6^, and contour^7,8^, and appear to be segregated according to some of these properties across the cortical surface^9^. Distinct from their purely sensory properties, V4 neurons are also known to encode information about behavioral and cognitive factors, particularly covert attention^10^, but also reward value^11^, and the direction of planned saccadic eye movements^12^–^14^. As with other neocortical areas, V4 is organized by a characteristic laminar structure, in which granular Layer 4 neurons receive feedforward sensory input from hierarchically ‘lower’ visual cortical areas, namely area V1 and V2^1,15^–^17^. Projections from area V4 to hierarchically ‘higher’ visual areas, such as TEO and posterior inferotemporal (IT) cortex, originate largely from layers II-III ^1,18^, whereas layer 5 neurons project back to V1 and V2 and subcortically to the superior colliculus^18^–^20^.

Recent studies have found laminar differences in attention-related modulation of neural activity. Buffalo et al., (2011)^21^ observed that changes in LFP power due to the deployment of covert attention differed between superficial and deep layers; gamma-band increases were found in superficial layers and alpha-band decreases were found in deep layers. Increases in firing rate with attention were observed to be similar in both laminar divisions. Nandy et al. (2017)^22^ compared attention-driven changes in spiking activity across three laminar compartments of V4 and observed significant firing rate modulation in superficial, granular and deep layers. In addition, they observed subtle, but reliable, differences in other aspects of activity across layers (e.g. spike count correlations). However, no previous studies have compared stimulus tuning properties, or looked for differences in other types of behavioral modulation across layers.

To investigate the layer dependence of stimulus and behavioral modulation in area V4, we measured the selectivity of V4 neurons to both factors in monkeys performing an attention-demanding task that dissociated covert attention from eye movement preparation. We then compared orientation tuning and behavioral modulation in superficial and deep layer individual units, and populations.

## Results and Discussion

Two monkeys (G and B) were trained to perform an attention-demanding task^23^ that required them to detect orientation changes in one of four peripheral oriented grating stimulus patches while maintaining central fixation (Figure 1a; see Experimental Procedures)^12^. Upon detection of a change, monkeys were rewarded for saccadic eye movements to the patch opposite the orientation change. Both monkeys performed well above chance. We recorded the activity of 698 units (277 single-units and 421 multi-units) at 421 sites using 16-channel linear array electrodes while monkeys performed the task. Electrodes were delivered perpendicular, or nearly perpendicular, to the cortical surface as guided by magnetic resonance imaging, and confirmed by receptive field (RF) alignment (Figure 1b). In each recording session, data from the 16 electrode channels were assigned laminar depths, relative to a common current source density (CSD) marker (Figure 1c, see Methods).

**Figure 1.**
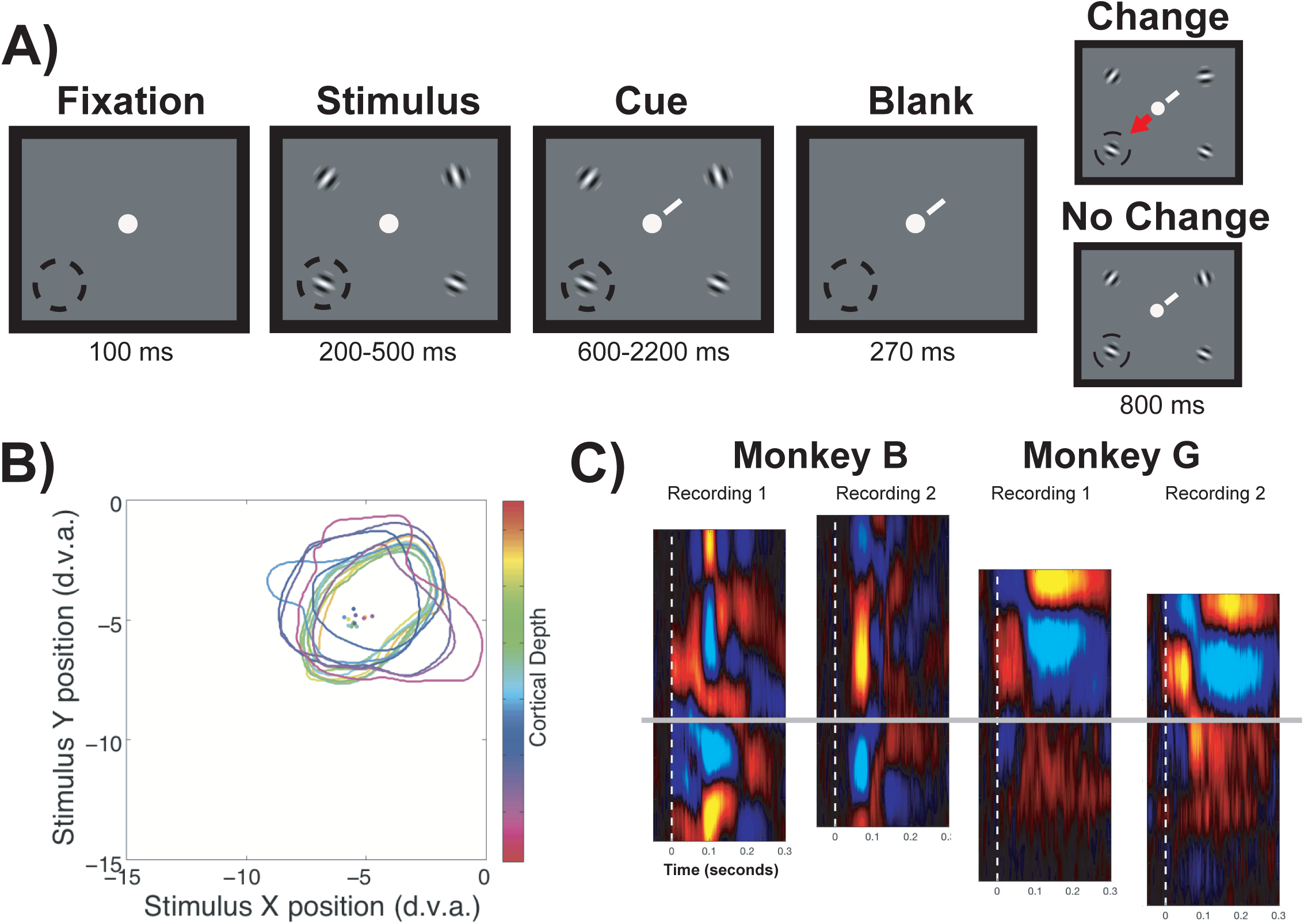
Behavioral task and perpendicular recordings in area V4. A) Panels depict phases of the attention task, and lower left dashed circle denote RF position of recorded neurons, and was not seen by subjects. Receptive fields of recorded neurons was always in the lower left, as indicated by the dashed circle outline. Task began with fixation at a central fixation point. Following fixation, randomly oriented Gabor gratings appeared at four positions. After an additional period, a cue (white diagonal line) appeared near the fixation point and indicated which grating was the target. A blank period followed in which the gratings disappeared, and then the stimuli reappeared on the screen with the target presented either at the same orientation or at a new orientation. Monkeys were rewarded for making saccadic eye movements to the stimulus opposite the changed target (arrow) or for maintaining fixation when the orientation did not change. B) Colored contours and corresponding dots respectively show the RF borders and RF centers mapped at electrode channels across difference cortical depths for an example V4 recording. C) Example current source density (CSD) with alignment feature for the two monkey subjects. The delineation between superficial and deep layers is indicated by the gray line.

### Orientation Selectivity

We first examined the proportion of units exhibiting significant orientation tuning and compared that proportion across layers (see Methods). Overall, 63.75% (445/698; P < 0.001) of units were significantly tuned for stimulus orientation (Figure 2a). Of these, we found that a significantly higher proportion of superficial units (74.9%) were tuned compared to deep units (58.3%; Chi-squared, P = 9.7×10^-6^). Next, we fit Gaussian functions to the normalized mean firing rates elicited by the eight orientations for each of the 698 units (Figure 2b, see Methods). Across superficial and deep layers, 35.5% (248) of units were well-fit (R^2^ > 0.7). Among the well-fit units, 98 were recorded in superficial layers (36.6% of superficial units) and 150 were recorded in deep layers (35% of deep neurons). These proportions were not significantly different from each other (Chi-squared, P > 0.05). Comparing fit parameters, we observed no significant differences in width or baseline between superficial and deep layers (width, superficial = 0.84, deep = 0. 67, P > 0.05; baseline, superficial = 0.10, deep = 0.10, P > 0.05). However, the mean amplitude of superficial layer units was significantly greater than that of deep layer units (superficial = 0.17; deep = 0.13; P = 0.0179).

**Figure 2.**
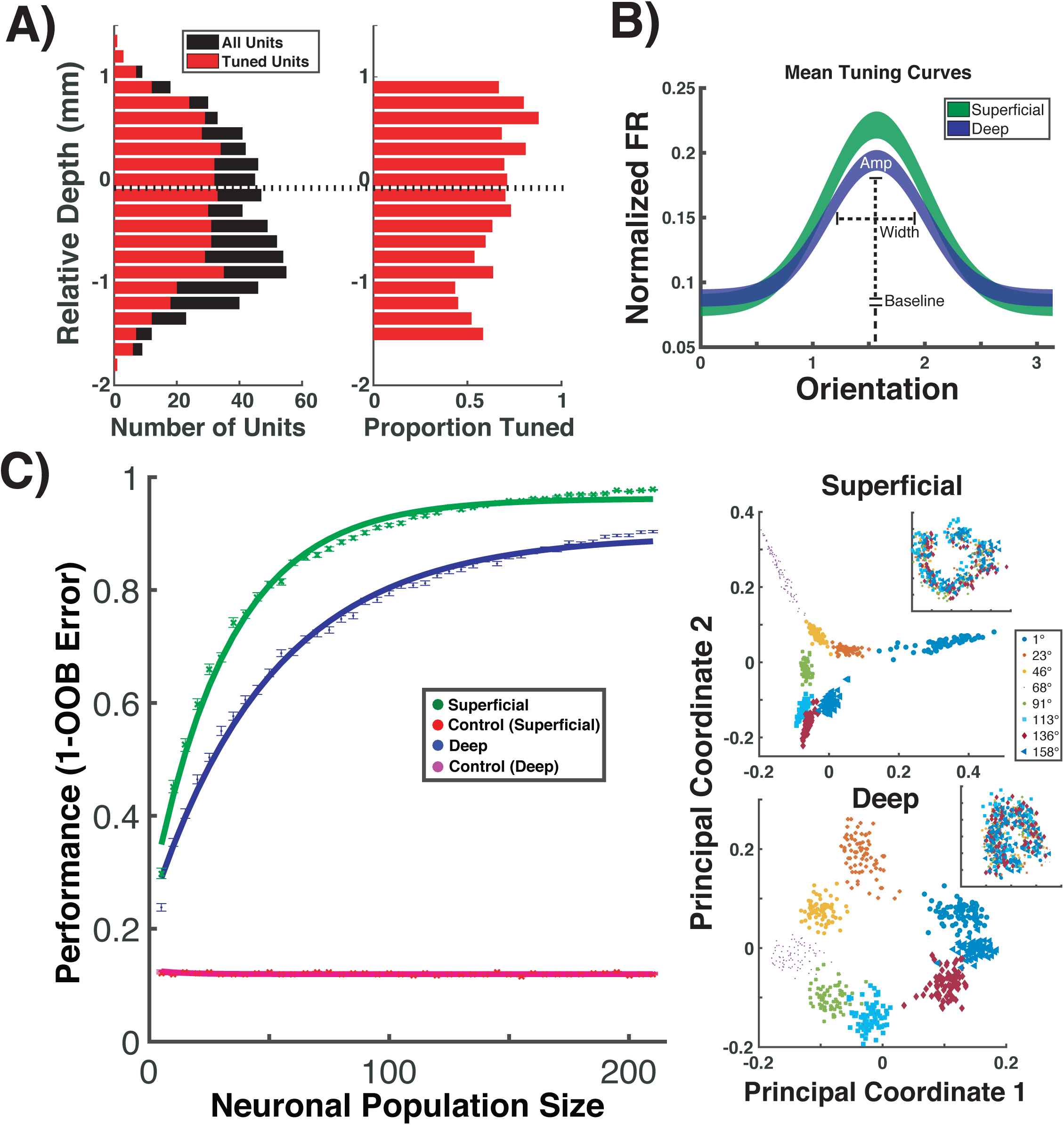
Orientation tuning in superficial and deep layers of area V4. A) Left, distribution of tuned units (red) among total units recorded (black) across cortical depth, relative to the superficial/deep CSD border. Right, the same data plotted as a proportion. B) Average Gaussian tuning fits, and definitions of fit parameters, for superficial (green) and deep (blue) neurons. Line thicknesses denote ±SEM. C) Left, performance of a Random Forest classifier at decoding stimulus orientation across different population sizes of superficial (green) and deep blue) neurons, along with shuffled controls for both (red and purple). Points indicate median values, and bars indicate the SEM for the 100 decoder cycles at each size. Solid lines indicate the fit saturating function. Right, multidimensional scaling (MDS) of classification for one cycle at the maximum population size (210 neurons). Each color/shape combination is associated with a unique orientation. Inset depicts the same MDS analysis after shuffling stimulus orientation labels.

Measurements of orientation tuning in individual units indicate that superficial layer units in our dataset were better tuned to stimulus orientation than their deep layer counterparts. However, we considered that these measurements might not capture all of the information conveyed about stimulus orientation. We therefore took a population decoding approach^24^ to measure the information available about orientation in the activity of all units within superficial or deep layers (see Methods). Decoder performance was computed as a function of neuronal population size. We then fit a “neuron-dropping” curves (NDCs)^25^ to the values, and compared the confidence intervals of the fit parameters for slope (b) and asymptote (c) for superficial and deep populations. Both superficial and deep units performed significantly above chance for all population sizes greater than zero. The NDS curve for superficial populations had a significantly greater slope (superficial b = 0.03071, 95% CI: 0.03002, 0.0314; deep b = 0.01976, 95% CI: 0.01925, 0.02026), and asymptotic performance was about 7% higher than deep units (superficial, c = 0.9622, 95% CI: 0.9597, 0.9647; deep, c = 0. 8969, 95% CI: 0.8931, 0.9008). Thus, as with the single-unit analysis, we found that stimulus orientation was more accurately encoded by populations of superficial layer neurons.

The robust differences in orientation selectivity we observed between the superficial and deep layer units raise important questions, such as whether those differences result simply from the known compartmentalization of orientation versus color tuning across V4^9^. However, even if we had oversampled one compartment or the other (e.g. more color compartments), doing so would not be expected to introduce an overall bias between upper and lower layers. It is also worth noting that since the primary evidence of feature-specific compartments in V4 comes from optical imaging, where much of the signal derives from superficial layers^26^, those compartments may be less well-defined within infragranular layers. Indeed, anatomical evidence indicates that intrinsic horizontal connections in V4, which appear to reciprocally connect columns across millimeters of cortex, exist predominantly in superficial layers, similar to earlier (e.g., V1, V2) and later stages of visual cortex^27^.

Second, our results raise the important question of whether the selectivity to other features, e.g. color or contour, is also greater in superficial layers. For example, substantial previous evidence suggests that neurons in V4 are unique in the computation of stimulus contour, not orientation, the former deriving from the orientation-specific input they receive from V1 and V2^7,8,28,29^. In such a case, our observations within orientation selectivity might not generalize to all other types of selectivity. Instead, the results might only generalize to features computed at earlier stages. Nonetheless, our results reveal the importance of assessing the laminar dependence of stimulus selectivity across visual cortex.

### Coding of Eye Movement Preparation and Covert Attention

We next examined activity across superficial and deep layers when monkeys covertly attended the visual stimulus, prepared a saccade to that stimulus, or ignored it. We first compared the average modulation for individual neuronal recordings made at varying laminar depths aligned to the superficial/deep boundary (Figure 3A). Overall, modulation across depth was significantly greater during eye movement preparation than during covert attention (P = 0.0024), a result we reported previously^10^. However, we observed no significant main effect of depth (P > 0.05), or an interaction of attention type and depth (P > 0.05). Nonetheless, movement-related modulation appeared to peak within the deep layers, suggesting that the difference in attention type was due to greater eye movement modulation in those layers. Thus, we directly compared the magnitude of modulation in the two attention types collapsed within superficial or deep layers. This revealed that while there was no significant difference in modulation in superficial layers (P > 0.05), saccade modulation was significantly greater within deep layers (P = 0.0041).

**Figure 3.**
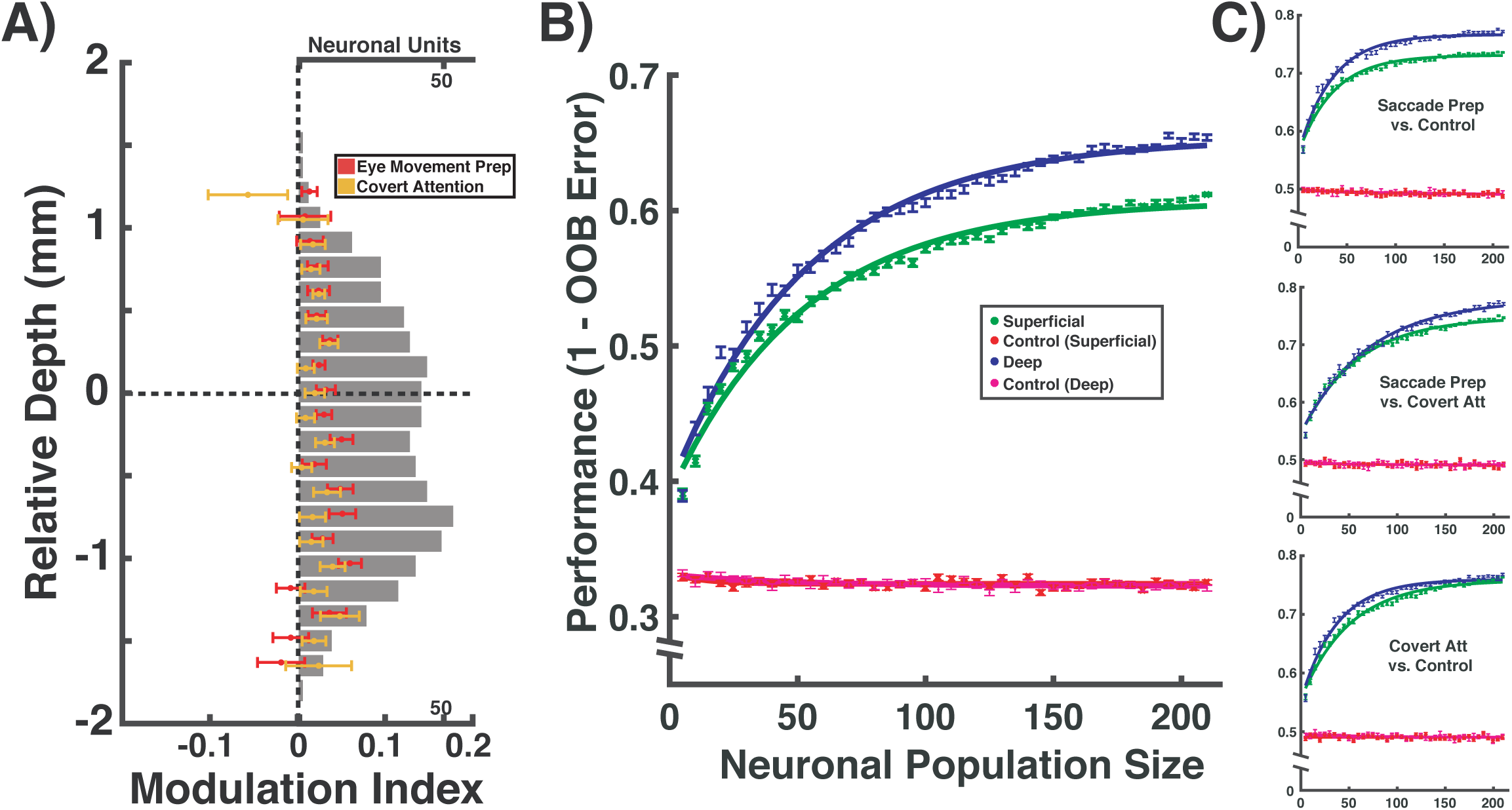
Behavioral modulation in superficial and deep layers of V4. A) Modulation indices across cortical depth. Individual medians and SEMs are plotted at each depth for covert attention (yellow) and saccade preparation (red), along with the total number of units recorded (grey). Depths with fewer than five neurons were removed. B) Performance of Random Forest decoder at distinguishing between the three behavioral conditions (covert attention, saccade preparation or control) from superficial and deep neurons, as a function of neuronal population size. C) Performance of the decoder at distinguishing between pairs of conditions: (top) saccade preparation from control; (middle) saccade preparation from covert attention; (bottom) covert attention from control.

Next, as with stimulus orientation, we decoded the behavioral condition using population activity from superficial (277 units) or deep (419 units) layers (Figure 3b), and classified activity as occurring during covert attention, saccade preparation, or control trials. The performance of decoding deep populations was significantly greater than superficial at all populations of >30 units. Although the slopes of the NDS fits were not significantly different, (superficial b = 0.01918, 95% CI: 0.01842, 0.01993; deep b = 0.01849, 95% CI: 0.01773, 0.01925), the asymptotic performance for deep units exceeded that of superficial units by more than 5% (superficial, c = 0.6073, 95% CI: 0.6053, 0.6092; deep, c = 0.6534, 95% CI: 0.6509, 0.6559). Thus, the behavioral condition was more accurately encoded by populations of deep layer units.

To investigate the conditions driving performance, we then conducted pairwise decoding of attentional conditions (Figure 3c). When decoding covert attention versus control, we found that although the NDC slope for deeper populations was greater than that of superficial populations, (superficial b = 0.01974, 95% CI: 0.01882, 0.02066; deep b = 0.02671, 95% CI: 0.0255, 0.02792), asymptotic performances were not significantly different (superficial, c = 0.7565, 95% CI: 0.7544, 0.7586; deep, c = 0.758, 95% CI: 0.7564, 0.7596). In decoding saccade preparation versus covert attention, we found a greater slope for superficial layer units, (superficial b = 0.01739, 95% CI: 0.01646, 0.01832; deep b = 0.01446, 95% CI: 0.01361, 0.01531), but a greater asymptotic performance for deep layer populations (superficial, c = 0.7475, 95% CI: 0.7448, 0.7503; deep, c = 0.7785, 95% CI: 0.7745, 0.7825). Lastly, when decoding saccade preparation versus control, we found that although the slopes were not significantly different, (superficial b = 0.02765, 95% CI: 0.02639, 0.02891; deep b = 0.0286, 95% CI: 0.02714, 0.03006), the asymptotic performance was ∼3% greater for deep units (superficial, c = 0.7295, 95% CI: 0.7282, 0.7308; deep, c = 0.7651, 95% CI: 0.7634, 0.7668). Thus, coding of attentional state, covert or overt, was greatest for units in the deep layers, where eye movement preparation was most strongly encoded.

Few studies have examined the influence of motor preparation on the responses of neurons in visual cortex, yet it is nonetheless clear that visually driven activity is affected by impending eye movements at many stages of the primate visual system^30^–^33^. Moreover, we have shown previously^12^, and in the present study, that the movement-related modulation of V4 activity is not only dissociable from modulation by covert attention, but it is more reliable. Those findings are consistent with the hypothesis that visual cortical areas contribute directly to visually guided saccades, particularly the refinement of saccadic plans according to features coded by particular visual areas (e.g. shape in area V4)^34^–^36^.

Our observation of stronger eye movement-related modulation in deep layers is also consistent with the fact that projections to the superior colliculus emanate principally from layer V pyramidal neurons throughout extrastriate visual cortex^37^. Moreover, deep layer neurons are a major source of feedback projections^1^, and thus the relative robustness of behavioral signals within deep layers may reflect the projection of those signals to earlier stages of visual processing. Consistent with this notion, a previous study of attentional effects in areas V1, V2 and V4 found evidence of a “backward” progression of modulation in these areas that begins in V4 and proceeds to V1^21^. Thus, the unique contributions of deep layer neurons to oculomotor output and in top-down influences may account for their superior coding of behavioral variables.

## Conclusion

We observed significantly greater orientation selectivity among units within the superficial layers of V4 using both tuning measures in single neurons and decoding of population activity. In contrast, using both single-unit and population activity, we observed that deeper layers conveyed more information about the behavioral relevance of visual stimuli. In particular, we found that neurons within deep layers conveyed more information than superficial neurons about the planning of saccadic eye movements. These results suggest a division of labor between superficial and deep layer neurons in the feedforward processing of stimulus features and the application of sensory information to behavior.

## Methods

### Subjects, Behavioral task, Visual Stimuli and Neuronal Recordings

Details of the subjects, the task, the stimuli and recording techniques are described in ^12^. In brief, two male rhesus macaques were surgically implanted with recording chambers. Monkeys were trained on an attention task that dissociated covert attention from saccade preparation. Trials were initiated when the monkey fixated a central point. After 100 ms of central fixation, a 300-500ms “stimulus epoch,” occurred, where four oriented Gabor patches appeared at four locations equidistant from the fixation point. This was followed by the “cue epoch,” lasting 600-2,200 ms. During this epoch, a line appeared near the central fixation point, directed toward one of the Gabor patches, indicating that it would potentially change orientations. After a variable interval, the array of stimuli disappeared briefly (270 ms) and then reappeared. Monkeys were trained to detect changes in orientation of any of the four stimuli upon reappearance. To dissociate the direction of covert attention from that of saccade preparation, monkeys were given a reward for responding to an orientation change with a saccade to the stimulus opposite the changed stimulus (i.e. antisaccade). If no change occurred at the cued location (50% of trials), the monkey was rewarded for maintaining fixation. Monkey G correctly responding on 69% of trials (77%, change trials; 62%, catch trials) and Monkey B correctly responding on 67% of trials, (62%, change trials; 70%, catch trials).

Electrophysiological recordings were made from area V4 on the surface of the prelunate gyrus with 16-channel, linear array U-Probes (Plexon, Inc., Dallas, TX). Electrodes were cylindrical in shape (180 mm diameter) with a row of 16 circular platinum/iridium electrical contacts (15 µm diameter) at 150 µm center-to-center spacing (total length of array = 2.25 mm). Recorded waveforms were classified as either “single neurons,” (277) or multi-neuron clusters (421). We use “units,” to refer to activity of both types.

## Cortical Column Laminar Recordings

### Electrode targeting: Use of MRI guidance to achieve perpendicularity

We sought to achieve simultaneous recordings at sites located within a single cortical column. In particular, the topographic organization of extrastriate visual cortex suggests that vertically separated neurons should have overlapping RFs, so we sought to record from a column principally by this definition. Since the cortical magnification factor (an estimate of how much cortical tissue corresponds to units of retinal space) is approximately 1 deg/mm ^38^, we could measure the approximate angle with the cortex by the distance between receptive fields measured on the deepest and most superficial recording contacts, and sought to keep this angle at 10 degrees or less, corresponding to a RF shift of ∼0.5 degrees, given 2 mm thickness of cortex.

In order to achieve these perpendicular penetrations we employed an MRI targeting technique ^39^. We implanted the monkeys with custom built recording chambers made from PEEK-type plastic, rather than from titanium, to avoid shadows in the MRI images. While we did not employ ceramic skull screws, we took some care to ensure that the titanium skull screws and plates were not located close to the recording chamber and brain areas of interest. We filled a custom-made plastic cylinder with copper sulfate solution. We anesthetized the monkey and inserted this cylinder into the recording chamber, into which it fit snugly. We performed structural MRI imaging (1.5 Tesla; T-1 weighted image) to visualize the location and orientation of the recording chamber (visible due to the high-contrast copper sulfate solution within it) relative to the position of the prelunate gyrus within the brain. By manually identifying the contours of the prelunate gyrus, we could compute perpendicular vectors to it and project these back to the level of the electrode stage, thus identifying which penetration approach vectors were likely to yield perpendicular penetrations.

### Achieving desired approach vectors

We employed a custom-built targeting device to angle and rotate the electrode into any desired orientation and position in three dimensions. The device consisted of a “double-eccentric” mechanism for positioning the electrode in the x-y plane of the well, a tilting mechanism, and a rotating mechanism. All four coordinates could be set with sub-millimeter precision using notches engraved in the device. The V4 recording chambers on both monkeys projected from the monkeys’ heads at an angle such that there was a unique point on the chamber’s perimeter at the lowest elevation. This point was identified computationally in the MRI images and was identified on the chamber itself by filling the chamber with saline solution until the liquid first contacted the lip of the chamber. With this point of alignment between the MRI images and the physical well, the exact X, Y, tilt, and rotation coordinates for an approach vector specified by the MRI images were geometrically determined.

### Electrode targeting: Assessing perpendicularity with RF overlap

RF positions and extents were estimated by computing the number of multi-unit spikes recorded on each channel in the 200ms period following stimulus onset for each of probe location in a RF-mapping task. During this task, subjects fixated a small (∼0.3 d.v.a.) white dot against a medium gray background. On each trial the six flash positions were selected from one of the rows of the grid in random order. A horizontally oriented grating was flashed for 50 ms at each position, with a 150-250ms variable delay between flashes. The flashes occurred at a total of 36 locations on a 6×6 grid with 3 d.v.a. spacing (total coverage 15×15 d.v.a.). If the subject maintained fixation within a 1.8 d.v.a. square window until after the sixth flash, he received a juice reward.

The upper right position of the grid was at the fovea such that only the lower left visual field was covered by the mapping. This 6×6 matrix of response counts was cubic spline interpolated to produce the full “RF map” and a 75%-of-max contour was determined, defining the RF border. The center of mass of the portion of the RF map within the RF border was defined as the RF center. This analysis was performed after recording RF-mapping task responses but before the change-detection task, so that a stimulus position could be chosen at a location that fell within the RF borders for all channels. If such a position was found, the recording was included in further analyses.

### Electrode targeting: Depth alignment

We lowered electrodes into the brain rapidly (∼25μm/sec) until one channel was in the cortex, based on visual examination of LFP and spiking activity being recorded concurrently. Then we advanced the electrode slowly (∼5μm/sec) until the uppermost electrode contact was near the point of entering the brain, being recorded during advancement. We withdrew the electrode 500μm to release compression of the brain caused by the electrode. During this brief withdrawal, no apparent change in the LFP or spiking activity was observed, confirming that this served to relax the cortex rather than change the position of the electrode relative to the brain. This manipulation qualitatively improved stability and recording quality. After reaching this position, the full-field flash task was run to assess the depth.

During the full-field flash task, monkeys fixated a small (∼0.3 d.v.a.) white dot against a black background. The monitor turned maximal white for one frame (∼8ms) then back to black. The flash occurred six times per trial with variable delays in the range of 150-250ms. If the monkey maintained fixation within a 1.8 d.v.a. square window until after the sixth flash, he received a juice reward. Approximately 30 trials, or 180 flashes, were completed per day. We computed the current source density (CSD) response to the full-field flashes. The CSD reflects the spatial and temporal position of current sources and sinks (i.e. where current flows into and out of the extracellular space, respectively) along the length of the electrode, given certain assumptions likely to be true for our recordings (Mitzdorf, 1985). The CSD can be computed discretely as the second spatial derivative of the LFP for each point in time, that is:

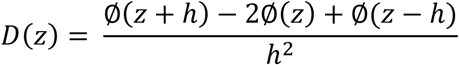

where z is the position in depth, h is the distance between potential measurements (in our case, 150μm), and *Φ* (z) is the potential measured as a function of depth. We also calculated the CSD according to the inverse estimation method ^40^, and display the results of this calculation, which produces smoother, higher resolution plots of CSD, in figures for clarity. However, results were qualitatively indistinguishable with both methods. Borders between current sinks of interest were manually identified and channel depths were computed, in mm, relative to these borders.

### Depth registration

In all included recordings, a prominent current sink was identified near the middle of the electrode, approximately 40-50ms after flash onset. This was often followed by another sink just below the first, peaking approximately 100ms after flash onset. These two sinks appeared in every included recording, and we therefore aligned the recordings on these functional markers of cortical laminae. In many recordings, further sinks were observed near the top of the probe at ∼150ms and near the bottom of the probe at ∼50ms. Because the widths of all four of these sinks, when present, were highly consistent from recording to recording, we assigned each channel a depth relative to this central feature.

Depths were measured in millimeters, and positive depths indicate channels superficial relative to the CSD feature. In some sessions, further CSD recordings at deeper locations revealed that no further current sources or sinks of comparable magnitude could be identified below these CSD features, assuring us that our electrode covered the depth of cortex. Two alignments of these functionally defined layers with anatomical cortical layers seem possible.

The uppermost sink could correspond to layer 2/3 (together), and the larger sink to layer 4 (Figure 1c). Alternately, the four visible sinks could correspond to layer 2, 3, 4, and 5 in order from superficial to deep. On the one hand, the first assignment seems reasonable as the thickness of the layers known histologically matches the thickness of these CSD features reasonably well, and our expectation from primary sensory areas is that layers 4 and 6 will have the earliest responses ^41^–^43^. However, the cortex may well be compressed around the electrode as it is inserted thus skewing the measured layer thicknesses. Layer 2 and 3 are well-differentiated cytoarchitecturally in V4 unlike in V1, suggesting they may not appear as a single sink. Furthermore, the earliest driving visual inputs into V4 are probably not from the ventral stream ^44^, which project into layer 4 ^15^, and may instead arrive from the pulvinar nucleus of the thalamus ^45,46^, which synapses into deep layer 3 ^47^ (Jones, 2007). This would indicate that the lower sink may correspond with the N95 marker used in previous studies to identify the granular layer ^42,48^–^50^.

## Data Analysis

### Tuning and Modulation Indices

To determine the tuning of each single neuron, we calculated the firing rate on each trial during a 300ms block, from 50ms to 350ms relative to stimulus onset. We then labeled the trials by stimulus orientation, and used a Kruskal-Wallis test to compare orientation distributions. If the p<0.001, we categorized the neuron as tuned. We then used a Chi-squared test to compare the proportion tuned in superficial versus deep layers. We also fit a Gaussian tuning function to the each neurons average firing rate for the eight stimuli using parameters for amplitude (a), preferred orientation (b), width (c) and baseline (d). The formula was given by:

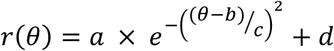

To obtain the parameters and goodness of fit measures, we used the Matlab fit function with nonlinear least squares, and constraints of 0 for the lower bound of all variables, and an upper bound of π for b and 8 for c. To determine if the neuron was well fit by the function, we used an adjusted R^2^ cutoff of 0.70. For each neuron, the averaged firing rates were rotated around π until the optimal fit was achieved. We then compared the function parameters of superficial and deep layer neurons. As the sample sizes of superficial and deep neurons were unequal, we used bootstrapping without replacement to match the sample sizes, and repeated each test 1000 times. The reported p-values are the mean of those produced by a Wilcox signed-rank test.

### Attention Modulation

For each neuronal unit, we calculated the mean firing rate during the cue epoch from −500ms to 0ms relative to the blank period. For each unit, we then calculated the attention modulation indices for eye movement preparation and covert attention relative to the orthogonal control, using the standard formula:

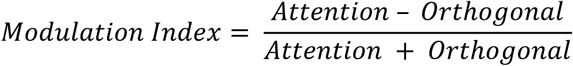

We then used a mixed effects model with fixed effects for neural depth, attention condition and an interaction term (implemented with the R package nlme^51^). To make layer comparisons within this omnibus model, we used three orthogonal contrasts: superficial attention conditions, deep attention conditions and superficial neuronal units versus deep neuronal units. In all tests, we included a random intercept for each neuronal unit, to control for repeat measures.

### Stimulus and Attention Classification

#### Feature Matrix

We assembled a dataset composed of neuronal firing rates recorded across the columnar arrays and across multiple experimental sessions (23 sessions from Monkey G; 20 sessions from Monkey B; 86 superficial neurons; 181 deep neurons) for all units for which we recorded a minimum number of trials per orientation (20), or attention condition (200). Each column of the feature matrix was a specific neuron’s firing rate, and each row of that column was the neuron’s firing rate on a specific trial. The rows of each column were aligned, so that they shared the same label for orientation or attention condition (depending on the epoch). The number of rows associated with each orientation or attention condition were matched, so that chance level was 12.5% for the orientation epoch and 33% for the cue epoch. Each neuronal unit had multiple columns in the feature matrix, corresponding to the number of bins in which firing rates were calculated. The firing rates for the orientation and cue epochs were calculated in two 150ms time bins, from 50ms following stimulus onset to 350ms following stimulus onset. This provided a gross temporal pattern which was noted to improve performance in Nandy et al. (2016)^24^. When building feature matrices with variable population sizes, we randomly sampled a population that size from all available units. This process was repeated 100 times, generating a unique of feature matrix for each run of the decoder.

#### Random Forest Classification

We used a Random Forest decoder, similar to that used in Nandy et al. (2016)^24^, as implemented by Matlabs (Mathworks TM) treebagger function. In addition to decoding based on firing rate, Random Forest can decode based on differences in firing rate variability, even when mean firing rates are equal^52,53^. Furthermore, rather than comparing each orientation to the others in turn, the decoder simultaneously considers all orientations. The decoder’s decision trees were trained on bags of trials (matrix rows), selected through bootstrapping with replacement, and tested each decision tree on trials not included in the training bag. This out-of-bag (OOB) error was used as the performance measure. It is significantly more conservative than cross validation, but has the advantage of using all available data when training the decoder. Furthermore, the bootstrapped sampling method has the traditional advantages associated with bootstrapping, such as revealing the true underlying distribution from the available training data, and reducing the impact of outlier trials ^53^. The decoder then used a boosting method to create decision trees. At each branch point, a random subset of the features (square root of the total number of features) was chosen to calculate potential decision boundaries. Each of the features in the subset was used as a linear threshold for linearly partitioning the population of trials. The Gini impurity (GI) of the original sample, as well as of the two partitions was calculated using the formula:

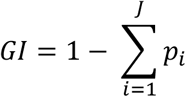

Where J is the number of classes, and *p*_*i*_ is the probability of choosing stimulus class *i* at random from the sample. The GI of the two partitions was averaged, and subtracted from the GI of the parent sample. The feature with the greatest decrease in GI was used at the decision boundary at that branch point. The use of a random subset of features reduces the influence of outlier features, allowing one to be less careful about the neurons selected for use in decoding. Stopping criteria for the decision trees was when either all the trials at a branch point had the same label (GI = 0), or there were only 5 trials at the branch point. We set the number of trees to 500. The decoder was trained and tested using each of the 100 feature matrices, producing a distribution of decoder performance.

For visualization, we calculated the proximity matrix based on shared decision leaves, and plotted the first two principal components for each trial.

#### Neuron-dropping Curves

We used neuron-dropping curves to assess the performance of the decoder. Also known as learning-curves, these are a standard tool in the machine learning to assess whether performance limitations are due to the decoder, or to the quantity of data. When computing these functions, the quantity of data used for decoding is varied and an error rate (or performance level) is plotted as a function of that quantity. The presence of an asymptote indicates that the decoder has reached maximal performance, whereas the absence of an asymptote indicates more data is needed. We then fit a saturating function and compared both the rate of rise, and the asymptotic value between populations.

We created pseudo-populations, starting with 5 units, and then incrementing by 5 until the maximal number of available units was reached. For each population size, we randomly sampled the requisite number from the larger population with replacement, repeating this process 100 times to bootstrap a representative distribution. To this range of performance levels, we fit the saturating function,

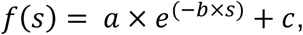

where s is the size of the population, a controls the y intercept, b the slope and c specifies the function asymptote. This was implemented using the Matlab fit function with the method non-linear least squares. A confidence interval of 95% was derived from the fitting process.

## Acknowledgements.

This work was supported by NEI grant EY014924 to T.M., a National Science Foundation graduate fellowship to N.A.S., and an HHMI medical research fellowship to W.W.P. We thank S. Hyde for valuable assistance with animal care, and B. Schneeveis for designing and building the 3D electrode angler.

